# Evaluation of Cancer Cell Membrane Disruption Property of Poly(vinyl alcohol)-ursodeoxycholic acid by 3D Cancer-Stromal Models with Macrophages

**DOI:** 10.64898/2025.12.22.696030

**Authors:** Kazuki Moroishi, Tova Hermodsson, Lennart Greiff, Malin Lindstedt, Michiya Matsusaki

## Abstract

In three-dimensional (3D) cancer tissue models, the low solubility of type I collagen (Col I) is a limitation in reproducing cancer tissue with high collagen density. we previously reported a “sedimentation culture method” using collagen microfibers (CMF) homogenized Col I powder. This 3D cancer tissue model is expected to reproduce tumor microenvironment comparable to that in tumors bearing mice owing to its high collagen density. However, this 3D cancer tissue model including immune cells such as macrophages has not been reported.

In this study, we fabricated a 3D culture system including normal human dermal fibroblasts, human umbilical vein endothelial cells, and cancer cells human monocytes cell line; THP-1 cells by sedimentation culture using CMF. 3D cancer tissue models co-cultured with human colorectal cancer cells for 5 days increased the expression levels of CD14 and CD206, markers of macrophages and M2 macrophages, in THP-1 cells, suggesting differentiation induction toward macrophages and polarization toward M2-like macrophages. We could also demonstrate the applicability of drug efficacy evaluation systems by evaluating the cytotoxicity of poly(vinyl alcohol)-ursodeoxycholic acid15 that induce cell death in response to weak acidic tumor microenvironment in 3D cancer tissue models with macrophages. This 3D cancer tissue models with macrophages are expected to be applied as tools for evaluating the efficacy of cancer therapeutic molecules and for understanding in detail the behavior of immune cells within the tumor microenvironment.

## 1. Introduction

Three-dimensional (3D) tissue culture is a technique that reproduces the microenvironment in tissues similar to that found in living tissue, while offering the advantage of high throughput as with 2D culture. 2D monolayer culture on rigid substrate could lead to genetic mutations in cells during 2D culture,^**1**^ whereas 3D culture systems such as spheroid and 3D printer would show different cell growth, morphology, differentiation, and cell-cell and cell-matrix interaction.^**2–5**^ In cancer cell treatment, it is important to construct a culture system that reproduces an environment similar to living tissue in order to properly investigate the delivery efficiency and efficacy.

3D scaffold materials that reproduce an extracellular matrix rich in collagen like a tumor tissue are required in cancer therapy. The ECM produced by fibroblasts is excessively accumulated, and collagen deposition and cross-linking are increased in accumulated ECM. This leads to the production of a cancer stroma with high elastic modulus, primarily composed of collagen I.^**6**,**7**^ Conventional 3D tumor tissue models (e.g., spheroids and collagen hydrogels) still struggle to reproduce tumor tissue with a high stromal density environment.

We previously reported “sedimentation culture” method using collagen microfibers (CMFs) that are micro-sized natural Col I with fiber lengths of 228 ± 147 μm and diameters of 13 ± 5 μm.^**8–12**^ CMFs enable the production of 3D tumor tissue models with high collagen density (up to 20–30% wt%) comparable to that in vivo by sedimentation culture, and can control elastic modulus up to more than 1.2 kPa. Furthermore, it is revealed that vascular endothelial cells and cancer stem cells in the 3D cancer tissue models with CMF showed vascularization, and high expression of cancer stem cell markers was observed, suggesting that they were highly malignant, similar to tumor tissue by CMFs.^**12**^ However, there is no report that include immune cells such as macrophages in sedimentation culture method by CMFs.

Immune cells in tumor stroma such as dendric cells (DCs), myeloid-derived suppressor cells (MDSCs), macrophages play an important role in regulating tumor progression.^**13**^ Macrophages differentiated from monocytes in blood in response to cytokine secreted in the tissue, and polarized into two major phenotypes: M1 macrophages and M2 macrophages. M1 macrophages are reported to have functions such as producing inflammatory cytokines and phagocytosing pathogens, while M2 macrophages are reported to promote tissue remodeling, suppress inflammation, and particularly promote tumor formation in tumor tissues.^**14**^ Therefore, cancer treatment that consider not only the cytotoxicity against cancer cells but also the surrounding immune cells and stroma are required. Although drug delivery system (DDS) is studied for a long time in cancer chemotherapy, the delivery efficiency of nanocarrier with anticancer agents *in vivo* experiment between tumor and cancer cells decreased 500-fold by injection dose due to uptake by macrophages.^**15**^ It is necessary to fully understand the behavior of macrophages in the tumor microenvironment to design molecules that adequately consider the presence of macrophages in cancer treatment.

In this study, we construct 3D culture system with CMF including normal human dermal fibroblasts (NHDFs), human umbilical vein endothelial cells (HUVECs), cancer cells, and THP-1 cells which human monocyte cell line (Figure 1a). Differentiation of THP-1 cells into macrophages was evaluated by co-culture with cancer cells. CD14 which is a macrophage marker in THP-1 cells upregulated by co-culture with colon cancer, suggesting differentiation into macrophages. Differentiated macrophages were also suggested to be polarizing toward M2-like macrophages. We also demonstrated cancer cell death induction property of PVA-U15 that show cytotoxicity in response to weak acidic tumor microenvironment in this 3D cancer tissue models (Figure 1b). PVA-U15 more significantly induced cancer cell death in weak acidic environment than neutral environment, demonstrating applicability of 3D cancer tissue models with CMF as an evaluation system of drug efficacy for cancer treatment. These results suggested that 3D cancer tissue models with CMF could reproduce tumor microenvironment. It is expected to be applied as a powerful tool for further understanding immune cells and cancer cells within the tumor microenvironment, and for providing effective cancer treatment approaches to individual patients by patient-derived cancer cells.

**Figure 1.**
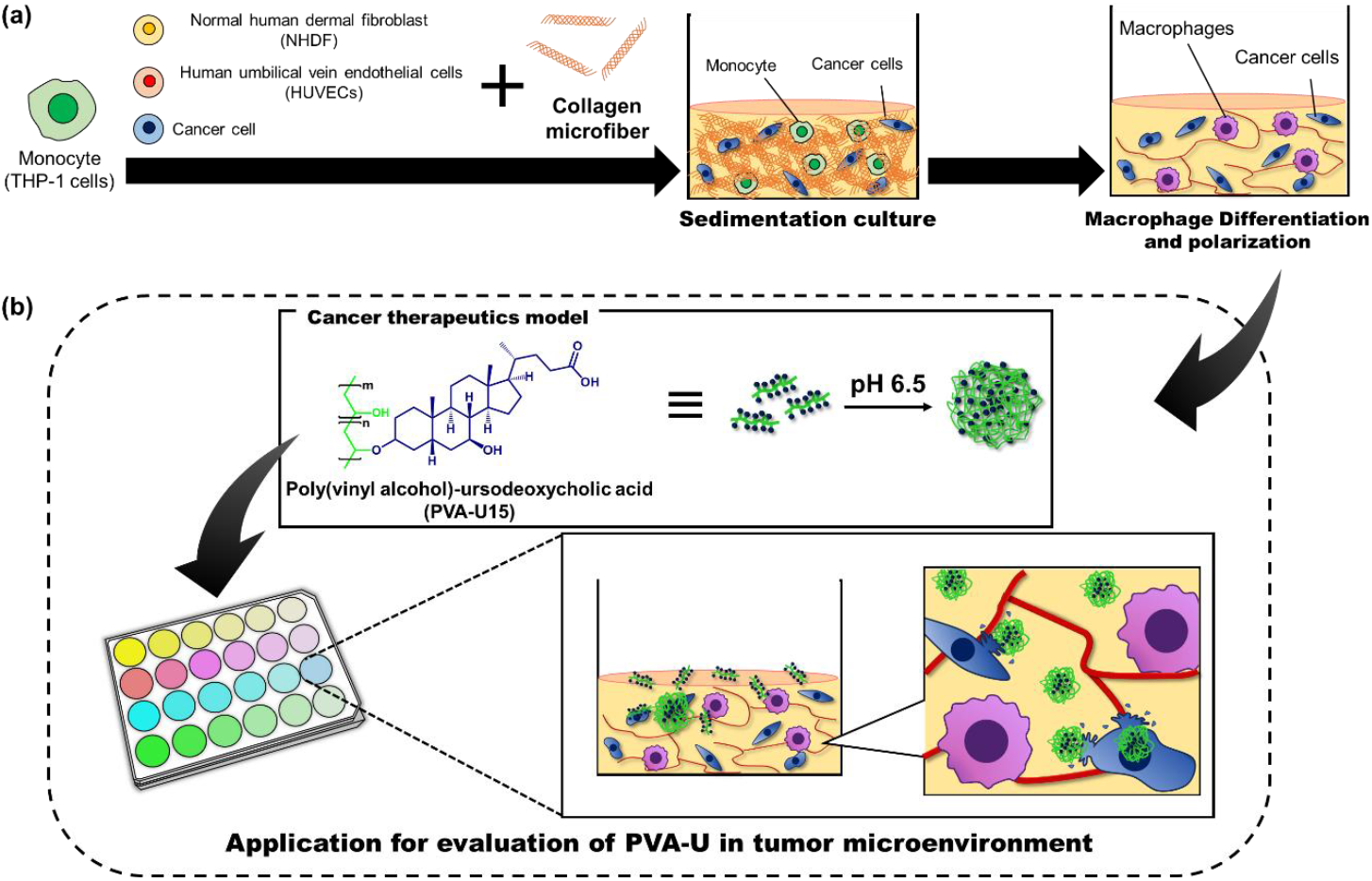
Overview of this study. (a)The three-dimensional cancer tissue model consisted of normal human dermal fibroblasts, human umbilical vein endothelial cells, cancer cells, monocytes, and collagen microfibrils homogenized from type I collagen powder as the extracellular matrix (ECM). These cells and materials were cultured after sedimentation by centrifugation. Monocytes cultured in such a high-stromal-density environment differentiate into macrophages and polarize into functional macrophages. (b) The schematic illustration of cell membrane disruption property evaluation of poly(vinyl alcohol)-ursodeoxycholic acid (PVA-U15) using 3D cancer tissue models with macrophages as a proof-of-concept. PVA-U15 is expected to disrupt cancer cell membrane by its weak acidic environment-responsive self-aggregation and cell membrane disruption property.

## 2. Result and discussion

### 2-1. Induction of differentiation of THP-1 cells into macrophages and polarization into M1 and M2 macrophages by co-culture with cancer cells in 3D tissue using collagen microfiber (CMF)

The 3D cancer tissue model with CMF is characterized by the formation of a vascular network driven by vascular endothelial cells under high stromal density. Therefore, as a preliminary study, we fabricated a 3D cancer tissue model using CMF by co-culturing normal human dermal fibroblasts (NHDFs), human umbilical vein endothelial cells (HUVECs), and pancreatic cancer (MiaPaCa-2) cells, and confirmed the formation of a vascular network after 5 days of incubation by immunostaining. These results showed a similar trend to previously reported 3D models with high mesenchymal density using CMF (Figure S1).^12^

This 3D cancer model was co-cultured with various cancer cell lines and THP-1 cells, a monocyte model, to evaluate whether THP-1 cells differentiate into macrophages. 3D cancer tissue models including THP-1 cells and colorectal cancer (HT-29) cells were similarly fabricated and cultured for 1 day and 5 days. Expression of CD14, CD86, and CD206, which are markers of macrophages, M1 macrophages, and M2 macrophages, respectively, was evaluated by immunostaining. The fluorescence of CD14 in 3D cancer tissue models incubated for 5 days was significantly higher than that of 1 day, suggesting differentiation of THP-1 cells into macrophages by co-culture with HT-29 cells (Figure 2a). Quantification of fluorescence intensity for CD14 in the 3D tumor tissue model after 5 days of incubation revealed a significant increase compared to that of 1 day (Figure 2b). In contrast, the expression of polarization markers showed that the fluorescence intensity of CD86, an M1 macrophage marker, did not change significantly after 5 days of incubation, whereas the fluorescence of CD206, an M2 macrophage marker, increased significantly after 5 days of incubation (Figures 2c, 2d). These results showed that THP-1 cells not only differentiated into macrophages but also induced polarization toward an M2-like phenotype by co-culture with HT-29 cells.

**Figure 2.**
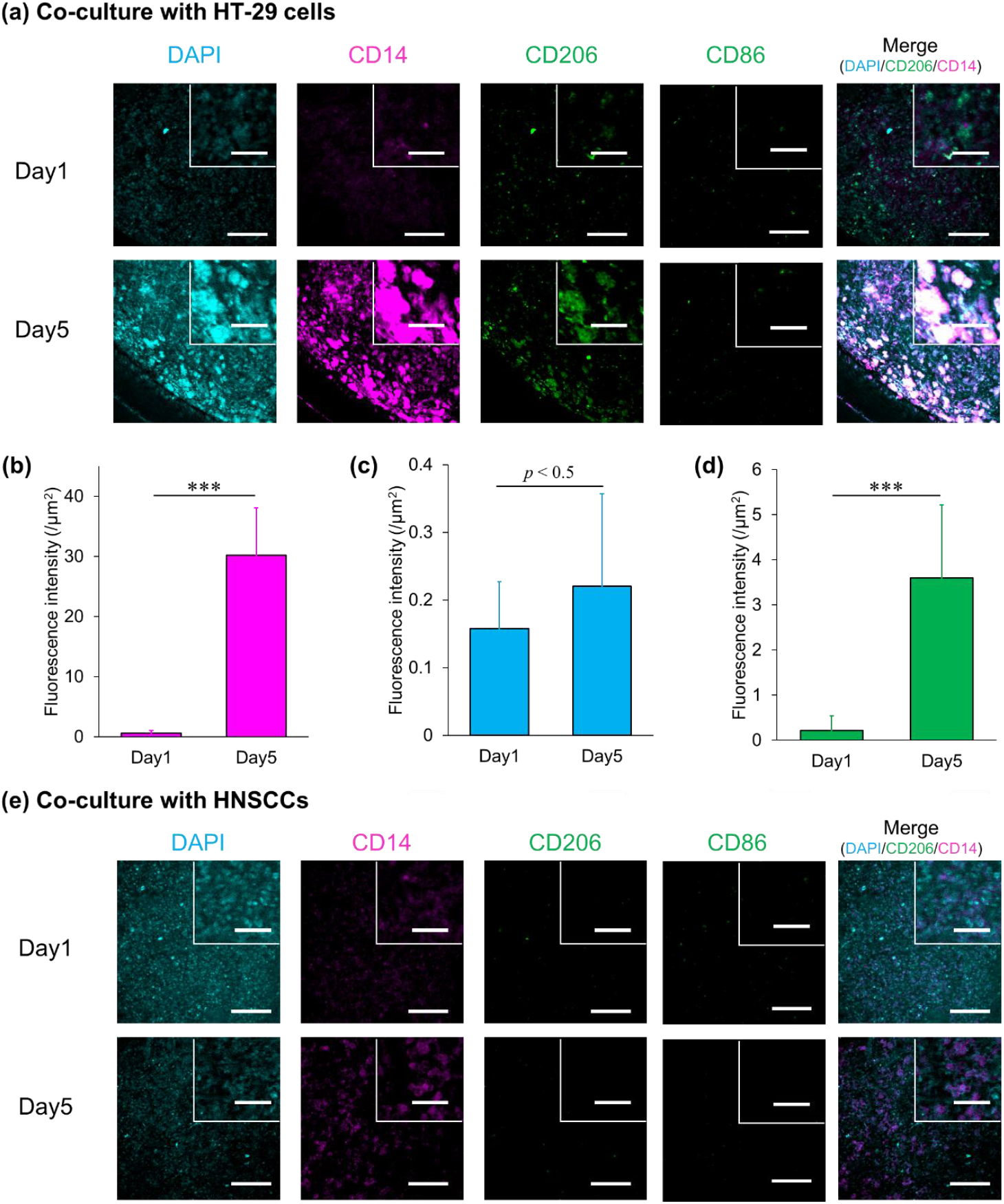
Differentiation of THP-1 cells in CMF tissue co-cultured with cancer cells. (a) Confocal images of 3D cancer tissue models with 3 mg of CMF co-cultured with HT-29 cells incubated for 1 day and 5 days stained with DAPI, CD14, CD86, and CD206. (b) Fluorescence intensity of CD14 antibody in 3D cancer tissue models with 3 mg of CMF co-cultured with HT-29 cells incubated for 1 day and 5 days. Fluorescence intensity of CD86 (c) and CD206 (d) co-localized with CD14 in 3D cancer tissue models with 3 mg of CMF co-cultured with HT-29 cells incubated for 1 day and 5 days. (e) Confocal images of 3D cancer tissue models with 3 mg of CMF co-cultured with HNSCCs incubated for 1 day and 5 days stained with DAPI, CD14, CD86, and CD206. Scale bars of image and enlarge images are 300 µm and 100 µm. Statistical analysis was performed using unpaired two-tailed Student’s *t*-test. Data are presented as mean ± S.D (\*\*\**p* < 0.001).

To confirm whether this differentiation and polarization depended on culture conditions, we evaluated the expression of CD14 and CD206 using a 2D culture system and a 3D model lacking NHDFs and HUVECs as control experiments.

Fluorescence of CD14 and CD206 was not detected in this model, confirming their expression is at very low levels (Figure S2). In contrast, expression of CD14 and CD206 was increased in a 3D cancer model mixed with NHDF, HUVEC, HT-29 cells, THP-1 cells, and 1 mg of CMF. These results suggested that the presence of CMF and both fibroblasts and vascular endothelial cells was important for the differentiation of THP-1 cells into macrophages and their polarization into M2-like macrophages.

The differentiation and polarization of THP-1 cells may be influenced by cytokines secreted from cancer and stromal cells, as well as the high-density collagen environment created by CMF.^16^ In particular, a high-density collagen environment may induce immunosuppressive properties in macrophages, promoting TAM-like behavior observed in the tumor microenvironment.^17^ Furthermore, it has been reported that fibroblasts promote the polarization of monocytes into M2-like macrophages via secreted factors, suggesting that NHDFs may have contributed to the differentiation and polarization of THP-1 cells in the 3D cancer tissue model.^18^

To investigate whether the differentiation and polarization of THP-1 cells demonstrated by co-culture with HT-29 cells was a phenomenon independent of specific cancer cells, we evaluated the induction of differentiation and polarization of THP-1 cells by co-culture with head and neck squamous cell carcinomas (HNSCCs). Tonsil cancer tissue, including HNSCC, has been reported to relatively abundant immune cells, such as macrophages, making it one of the important cancer types for analyzing immune cell responses in the tumor microenvironment^19^. In this study, HNSCC cells were co-cultured with THP-1 cells in a 3D cancer tissue model using CMF, and the induction of macrophage differentiation and polarization was evaluated. The 3D tumor tissue model co-cultured with HNSCC cells showed a significant increase in CD14 fluorescence intensity after 5 days of incubation compared to day 1, suggesting that THP-1 cells may have differentiated into macrophages (Figure 2e). In contrast, fluorescence for CD86 and CD206 was not detected at any culture day, and distinct M1-like or M2-like polarization was not observed (Table S1). These results suggested that the 3D cancer model using CMF co-cultured with HT-29 cells and HNSCCs induces macrophage differentiation in THP-1 cells, while polarization into M2-like macrophages may depend on the type of cancer cells and culture conditions. This model could provide a useful experimental platform for evaluating differences in macrophage responses to specific cancer cell types under conditions that replicate the tumor microenvironment.

### 2-2. Evaluation of cell death induction by poly(vinyl alcohol)–ursodeoxycholic acid (PVA-U15) using a macrophage-containing 3D cancer tissue model

The applicability of a 3D cancer tissue model derived from THP-1 cells differentiated into macrophages as an anticancer drug efficacy evaluation system was assessed. PVA-U15, which has been reported to self-aggregate and disrupt cell membranes in response to the weakly acidic tumor microenvironment, was used as a therapeutic molecule.

3D cancer tissue models with NHDFs, HUVECs, HT-29 cells, THP-1 cells, and 3 mg of CMF were cultured for 5 days, inducing differentiation of the THP-1 cells into macrophages. These models were treated with 100 µg mL^−1^ PVA-U15 for 24 hours at both pH 7.4 and pH 6.5 (Figure 3a). 3D cancer tissue models treated with PVA-U15 were observed using immunostaining with anti-EpCAM antibody and anti-Caspase-3 antibody (non-cleaved form), which are markers for HT-29 cells and dead cells, respectively.

**Figure 3.**
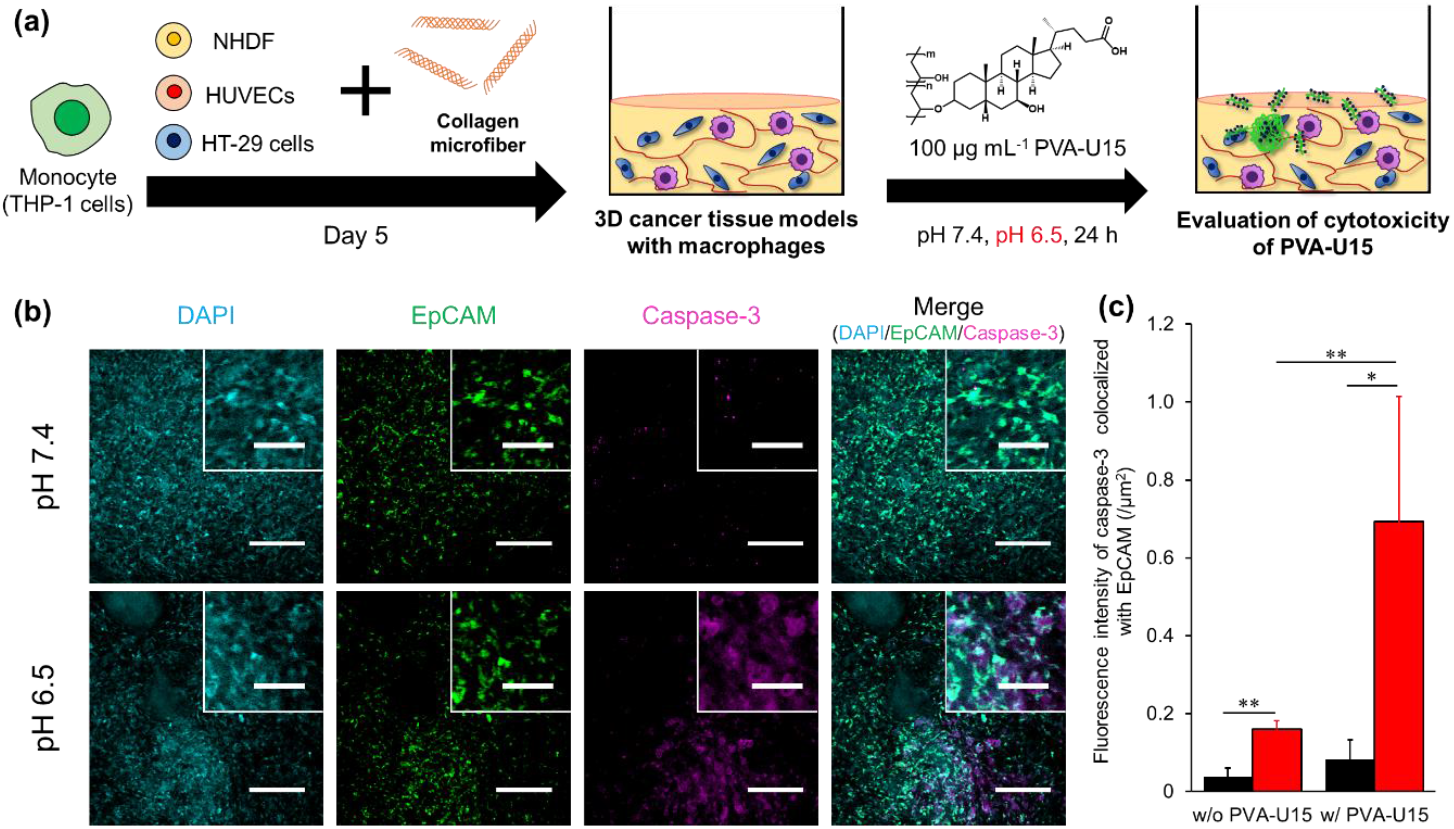
Evaluation of cytotoxicity of PVA-U using 3D cancer tissue model with macrophages. (a) Schematic illustration of the evaluation system for weak acidic environment-responsive cytotoxicity by PVA-U15. (b) Confocal images of CMF tissue treated with PVA-U15 at pH 7.4 and pH 6.5 stained with DAPI, EpCAM, and Caspase-3. Scale bars of image and enlarge images are 300 µm and 100 µm. (c) Fluorescence intensity of caspase-3 localized with EpCAM in CMF tissue treated with and without PVA-U15 at pH 7.4 (black) and pH 6.5 (red) (n = 3). Statistical analysis was performed using unpaired two-tailed Student’s *t*-test. Data are presented as mean ± S.D (\**p* < 0.05, \*\**p* < 0.01).

The fluorescence of Caspase-3 in the 3D cancer tissue model at pH 6.5 slightly increased compared to that at pH 7.4 (Figure S3). In contrast, the fluorescence of Caspase-3 in the 3D cancer tissue model treated with PVA-U15 at pH 6.5 markedly increased compared to that at pH 7.4, revealing that PVA-U15 induces cell death in response to a weakly acidic environment (Figure 3b). Co-localization between the fluorescence intensity of Caspase-3 and the fluorescence area of EpCAM was also quantified from confocal images (Figure 3c). The fluorescence intensity of caspase-3 increased 4.4-fold with PVA-U15 treatment at pH 6.5, demonstrating the induction of cancer cell death by PVA-U15 in a 3D cancer model containing macrophages.

Additionally, fluorescence of Caspase-3 was observed in areas that did not co-localize with EpCAM. Since simultaneous immunostaining was not performed for other cell types, such as macrophages, it was not possible to determine whether apoptosis occurred in these cells. However, these results suggested that cell death may also occur in cells other than cancer cells within 3D cancer tissue models, such as macrophages derived from THP-1 cells or NHDF, due to weak acidic environment-responsive cell membrane disruption of PVA-U15.

## 3. Conclusion

In summary, we fabricated a 3D cancer tissue model with high collagen density containing monocytes mixed with CMF, NHDF, HUVEC, cancer cells, and THP-1 cells. THP-1 cells differentiated into macrophages and were further induced to polarize into M2-like macrophages under co-culture with HT-29 cells and other stromal cells. Additionally, in 3D cancer tissue models containing macrophages, PVA-U15 selectively induced cancer cell death under weakly acidic conditions as previously reported, demonstrating its applicability as a three-dimensional cancer model. However, it should be noted that this study analyzed polarization from macrophages to M1-like and M2-like macrophages using only a single marker for each phenotype. Therefore, more detailed phenotypic analysis using other polarization-related markers present in each phenotype of macrophages is necessary in the future. These findings suggest that the 3D cancer model containing immune cells based on the CMF developed in this study is expected to provide a useful platform for evaluating the efficacy of cancer therapeutic molecules and analyzing immune cell behavior as an in vitro evaluation system that more accurately reproduces the tumor microenvironment.

## 4. Experimental Section/Methods

### 4.1 Materials

. A phosphate-buffered saline (PBS) was purchased from Nacalai Tesque (Kyoto, Japan). A Triton^TM^ X-100 and a bovine serum albumin (BSA) were purchased from Sigma-Aldrich (St. Louis, USA). A type I collagen derived from Porcine skin was purchased from nippi (Tokyo, Japan). 4% Paraformaldehyde Phosphate Buffer Solution was purchased from FIJIFILM Wako (Osaka, Japan). Tissue-clearing reagent CUBIC-R+(M) for animals was purchase from Tokyo Chemical Industry Co. Ltd. (Tokyo, Japan). A mouse anti-CD14 monocloal antibody (ref. ab181470), a rabbit anti-CD206 polyclonal antibody (ref. ab64693), an rabbit anti-CD86 monoclonal antibody (ref. ab239075), and a rabbit anti-EpCAM monoclonal antibody (ref. ab213500) were purchased from Abcam (Cambridge, UK). A mouse anti-Caspase-3 monoclonal antibody was purchased from Proteintech (Illinois, USA). A mouse anti-CD14 monoclonal antibody was purchased from Agilent Technology (California, USA). Goat serum, 4’,6-diamidino-2-phenylindole (DAPI), Alexa Fluor 647-conjugated goat anti-mouse IgG, and Alexa Fluor 647-conjugated goat anti-rabbit IgG were purchased from Thermo Fisher Scientific (Waltham, USA).

### 4.2 Cell culture

Head and neck squamous cell carcinoma (HNSCC) line, named LU-HNSCC-7, established in Lund University.^20^ Normal human dermal fibroblasts (NHDFs), human colorectal cancer cell lines HT-29, and HNSCCs were maintained in Dulbecco’s Modified Eagle Medium (DMEM, Nacalai Tesque, Kyoto, Japan) supplemented with 10% fetal bovine serum (FBS, Thermo Fisher Scientific, Waltham, USA). Human umbilical vein endothelial cells (HUVECs) were maintained in KBM VEC-1 Kohijn Bio, Saitama, Japan) supplemented with 500 µL of KBM VEC-1 supplement and 2% FBS. THP-1 cells were maintained in RPMI-1640 (FUJIFILM Wako, Osaka, Japan) supplemented with 10% FBS. Cell cultivation was performed at 37 °C in a humidified atmosphere of 5% CO_2_.

### 4.3 Differentiation of THP-1 cells co-cultured with cancer cells in a 3D cancer tissue model using CMF

30 mg of sterilized pieces of collagen sponge were homogenized for 1 min in 5 mL of 1× phosphate buffered saline (PBS) using a homogenizer (≈22000 rpm), and take 1 mg or 3 mg of CMF into each eppendolf tube. Each CMF was centrifuged at 10000 rpm for 1 min. 1 mg or 3 mg of CMF was used for each sample, NHDFs (1.0 × 10^6^), HUVECs (3.0 × 10^5^), THP-1 cells (2.0 × 10^5^), and cancer cells (MiaPaCa-2, HT-29, or HNSCC; 1.0 × 10^5^) were mixed with CMF in 200 µL of DMEM and transferred to 24-well transwell inserts. Each well was centrifuged at 1,100 g for 15 min and incubated for 1 and 5 days in 12 mL/well of DMEM supplemented with 10% FBS using a 6-well plate and self-made adaptors. As control experiments, THP-1 cells were cultured in 2D conditions or in 3D cancer tissue models without NHDFs and HUVECs. The medium was changed every 2-3 days. 3D cancer tissue models were fixed with 2.5 mL of 4% PFA Samples were incubated with 100 µL of mouse anti-CD31 antibody for evaluation of formation of vascular network, and mouse anti-CD14 antibody and rabbit anti-CD206 antibody or rabbit anti-CD86 antibody, respectively, for evaluation of differeintiation and polarization at 4 °C for 1 day. Samples were stained with Alexa Fluor 647-conjugated goat anti-mouse IgG, Alexa Fluor 647-conjugated goat anti-rabbit IgG, and DAPI for 3 hours. Samples were cleared using CUBIC reagent Immunostaining was similarly performed on 2D-cultured THP-1 cells, with each step involving centrifugation at 3500 rpm for 1 minute and resuspension in each solution. Samples for evaluation of vascular network were observed with confocal microscope (CellVoyager CQ-1, Yokogawa Electric Corporation, Tokyo, Japan). Other samples were observed with confocal laser scanning microscopy (FV10i, Olympus, Tokyo, Japan). Fluorescence intensity of anti-CD14, anti-CD206, and anti-CD86 were analyzed by randomly selecting 5 areas in the sample using ImageJ.

### 4.4 Application of 3D cancer tissue model using CMF for cytotoxicity evaluation of Poly(vinyl alcohol)-Ursodeoxycholi acid15 (PVA-U15)

A PVA-U15, which a PVA modified with UDCA at a grafting degree of 15%, was used as a proof-of-concept model to verify its applicability for evaluating drug efficacy in this 3D cancer tissue model. A PVA-U15 was synthesized as described in previous research reports. 30 mg of sterilized pieces of collagen sponge were homogenized for 1 min in 5 mL of 1× phosphate buffered saline (PBS) using a homogenizer at approximately 22000 rpm, and take 3 mg of CMF into each Eppendolf tube. Each CMF was centrifuged at 10000 rpm for 1 min. 1.0 × 10^6^ NHDFs, 3 × 10^5^ HUVECs, 1.0 × 10^5^ HT-29 cells, 2.0 × 10^5^ THP-1 cells, and 3 mg of CMF removed PBS were mixed with 200 µL of DMEM, and put into 24-well plate transwells. Each well was centrifuged at 1,100 g for 15 min and cultivated for 5 days in 12 mL/well of DMEM supplemented with 10% FBS using a 6-well plate and self-made adaptors. The medium was changed every 2-3 days. 1.2 mg of PVA-U15 were dissolved in 20 µL of DMSO to prepare each PVA-U solution at a concentration of 60 mg mL^-1^. These were diluted to 20 mg mL^-1^ with DMSO and 25 µL of these solutions were dissolved in 5 mL of DMEM supplemented with 10% FBS at pH 7.4 and 6.5, respectively to prepare polymer solutions containing 0.5% DMSO at concentrations of 100 µg mL^-1^. The medium was removed and each sample was transferred to a 24-well plate after 5 days of culture. Two samples were treated with 2.5 mL of 100 µg/mL PVA-U15 mixed medium and normal medium under conditions of pH 7.4 and pH 6.5. All samples were fixed with 2.5 mL of 4% PFA. Samples were incubated with 100 µL of mouse anti-Caspase-3 antibody and rabbit anti-EpCAM antibody, respectively, at 4 °C for 1 day. Samples were stained with Alexa Fluor 647-conjugated goat anti-mouse IgG, Alexa Fluor 647-conjugated goat anti-rabbit IgG, and DAPI for 3 hours. Samples were rendered transparent by adding 100 µL of CUBIC-R+(M) for animal, which was replaced with a fresh solution after 30 min and incubated at room temperature for 1 day. All samples were observed with FV10. Fluorescence intensity of anti-Caspase-3, anti-EpCAM were analyzed by randomly selecting 3 areas in the sample using ImageJ.

## Author contribution

K.M. carried out the experiments. K.M. wrote the manuscript. T.H., L.G., and M.L. provide head and neck squamous cell carcinomas K.M., M.L. and M.M. edited the manuscript. K.M., T.H., M.L. and M.M. contributed to the design and implementation of the research and the analysis of results. All authors have seen and approved the final manuscript.

## Acknowledgements

This work was supported by a JST SPRING (JPMJSP2138) from Japan Science and Technology Agency (JST). This study was financially supported by KAKENHI JP25KJ1752, JP22H05131, JP22H05138, JP22H05140, JP22H05141, JP25H01220, JP22K21348, and JP21H04634 from Japan Society for Promotion of Science (JSPS). This study was also supported by JPJSBP120239201, JPJSBP120252301, and Y2024L0906033 from JSPS, COI-NEXT (JPMJPF2009) from JST.

