## Supplemental information for "Evaluation of Cancer Cell Membrane Disruption Property of Poly(vinyl alcohol)-ursodeoxycholic acid by 3D Cancer-Stromal Models with Macrophages"

Supporting Information

### Differentiation of THP-1 Cells in 3D Cancer-Stromal Tissue by Collagen Microfiber for Replicating of Tumor Microenvironment

*Kazuki Moroishi[1], Tova Hermodsson[2], Lennart Greiff[3,4], Malin Lindstedt[2], and Michiya Matsusaki*[1]*

[1]Division of Applied Chemistry, Graduate School of Engineering, The university of Osaka, Osaka, Japan.

[2]Department of Immunotechnology, Lund university, Lund, Sweden

[3]Department of ORL, Head & Neck Surgery, Skåne University Hospital, Lund, Sweden,

[4]Department of Clinical Sciences, Lund University, Lund, Sweden,

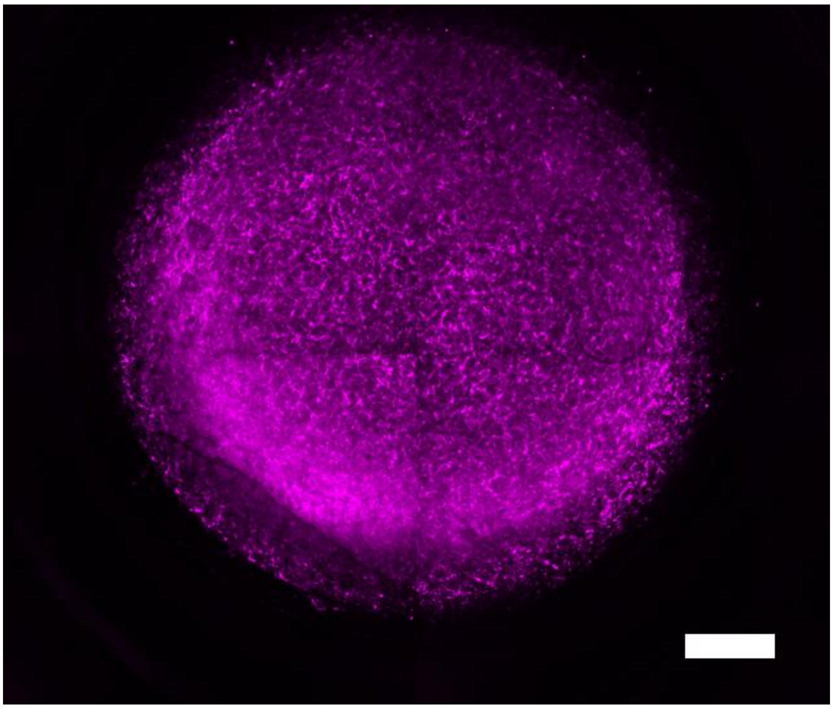

**Figure S1.** Confocal images of 3D cancer tissue models with 3 mg of CMF co-cultured with NHDFs, HUVECs, and MiaPaCa-2 cells These models were stained with anti-CD31 antibody. A scale bar of image is 1000 μm.

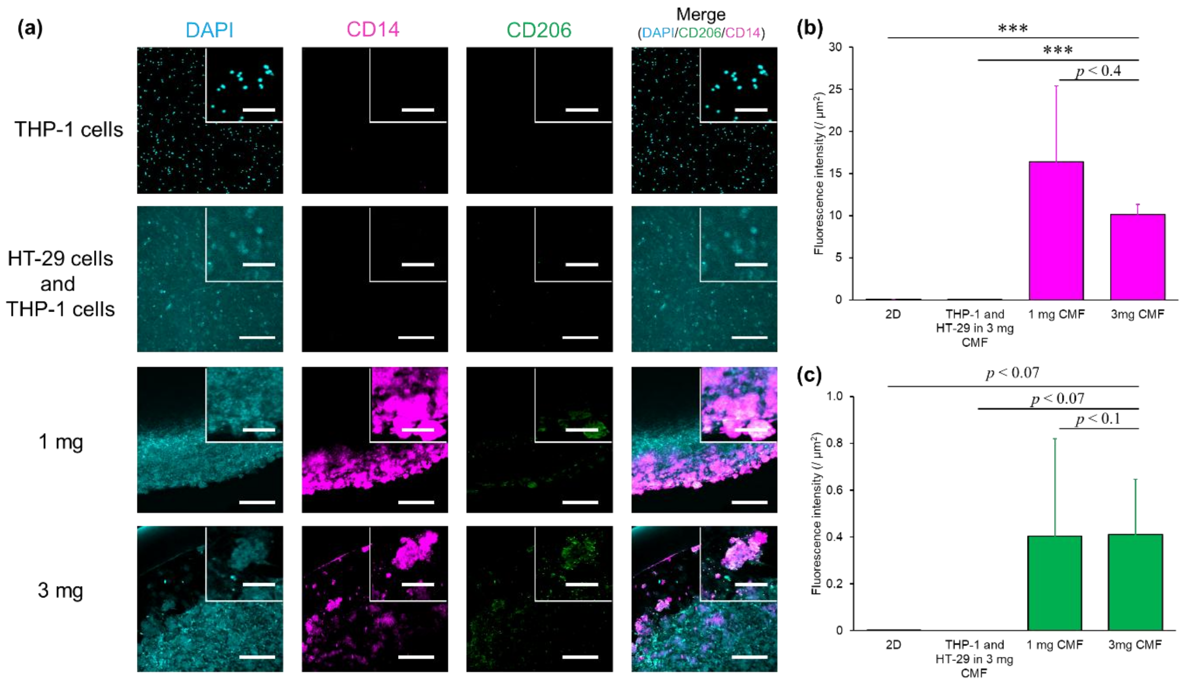


**Figure S2. Differentiation of THP-1 cells in 3D cancer models with modified stromal cells and collagen amount.** (a) Confocal images of THP-1 cells by 2D cultures, co-culturing with HT-29 cells alone in 3D cancer tissue model using 3 mg CMF, and co-culturing with NHDFs, HUVECs, HT-29 cells in 3D cancer tissue model using 1 mg and 3 mg of CMF incubated for 5 days, respectively. These models were stained with DAPI, anti-CD14 antibody, and anti-CD206 antibody, respectively. Scale bars of image and enlarge images are 300 μm and 100 μm. Fluorescence intensity of CD14 (b) and CD206 (c) in 2D culture, 3D cancer tissue model using 3 mg CMF co-cultured with HT-29 cells alone, and 3D cancer tissue model using 1 mg and 3 mg of CMF co-cultured with NHDFs, HUVECs, HT-29 cells incubated for 5 days, respectively. Statistical analysis was performed using unpaired two-tailed Student's t-test. Data are presented as mean ± S.D (***$p < 0.001$).

**Table S1.** Fluorescence intensity of CD14, CD86, and CD206 in 3D cancer tissue model using 3 mg of CMF co-cultured with HNSCCs.

| Merker | Day1 | Day5 |
|---|---|---|
| CD14 | 0.020 ± 0.020 | 4.6 ± 3.9 |
| CD86 | *N.D.* | *N.D.* |
| CD206 | *N.D.* | *N.D.* |

*N.D.*: not detected; fluorescence signals were below the threshold defined by secondary antibody–only controls.

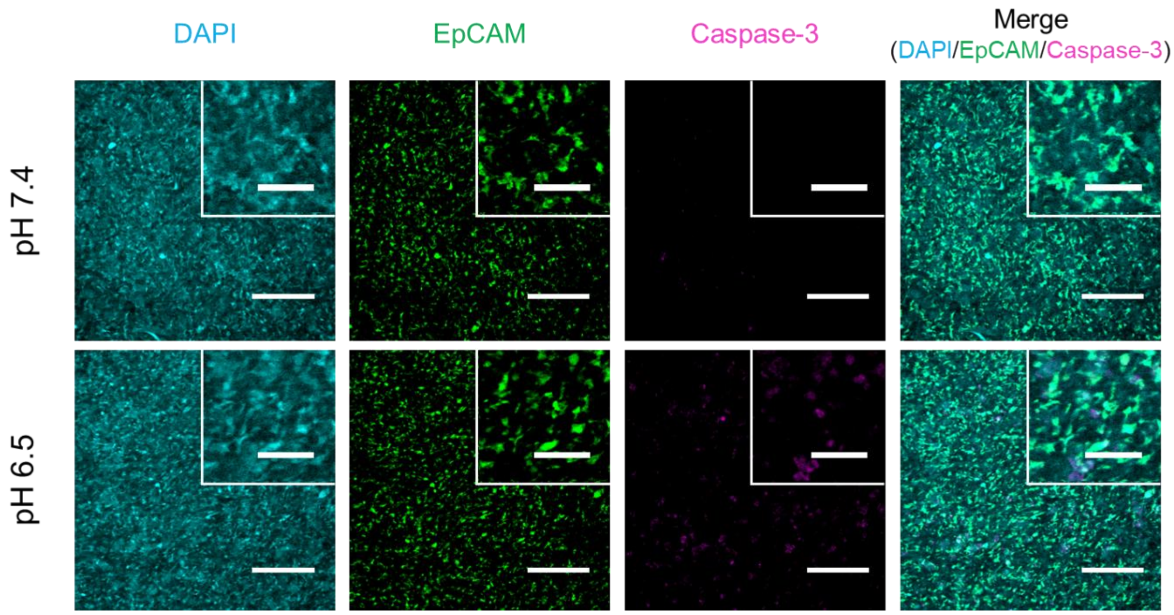


**Figurer S3.** Confocal images of 3D cancer tissue model at pH 7.4 and pH 6.5 stained with DAPI, anti-EpCAM antibody, and anti-Caspase-3 antibody. Scale bars of image and enlarge images are 300 μm and 100 μm.
